# Activation of IP3R in atrial cardiomyocytes leads to generation of cytosolic cAMP

**DOI:** 10.1101/2024.03.28.583721

**Authors:** Emily C Akerman, Matthew J. Read, Samuel J. Bose, Andreas Koschinski, Rebecca A. Capel, Ying-Chi Chao, Milda Folkmanaite, Svenja Hester, Roman Fischer, Thamali Ayagama, Steven D. Broadbent, Rufaida Ahamed, Jillian N. Simon, Derek A. Terrar, Manuela Zaccolo, Rebecca A. B. Burton

**Author notes:** ^†^These authors contributed equally to this manuscript.

## Abstract

Atrial fibrillation (AF) is the most common sustained cardiac arrhythmia. Excessive stimulation of the IP3 signaling pathway has been linked to AF through abnormal calcium handling. However, little is known about the mechanisms involved in this process. We expressed Fluorescence resonance energy transfer (FRET) based cytosolic cAMP sensor EPAC-S^H187^ in neonatal rat atrial myocytes (NRAMs) and neonatal rat ventricular myocytes (NRVMs). In NRAMs, addition of the alpha (α)-1 agonist phenylephrine (PE, 3µM) resulted in a bi-phasic FRET change (R1) 21.20 ± 7.43% and (R2) 9.67 ± 4.23% and addition of membrane permeant IP3 derivative, 2,3,6-tri-O-Butyryl-myo-IP3(1,4,5)-hexakis(acetoxymethyl)ester (IP3-AM, 20μM) resulted in a peak of 20.31 ± 6.74%. These FRET changes imply an increase in cAMP. Prior application of IP3 receptor (IP3R) inhibitors 2-Aminoethyl diphenylborinate (2-APB, 2.5μM) or Xestospongin-C (0.3μM) significantly inhibited the change in FRET in NRAMs in response to PE. Xestospongin-C (0.3μM) significantly inhibited the change in FRET in NRAMs in response to IP3-AM. The FRET change in response to PE in NRVMs were not inhibited by 2-APB or Xestospongin-C. Finally, the localisation of cAMP signals was tested by expressing the FRET-based cAMP sensor, AKAP79-CUTie, which targets the intracellular surface of the plasmalemma. We found in NRAMs that PE led to FRET change corresponding to an increase in cAMP that was inhibited by 2-APB and Xestospongin C. This data support further investigation of the pro-arrhythmic nature and components of IP3 induced cAMP signalling to identify potential pharmacological targets.

**NEW & NOTEWORTHY:** This study shows that indirect activation of the IP3 pathway in atrial myocytes using phenylephrine and direct activation using IP3-AM leads to an increase in cAMP and is in-part localized to the cell membrane. These changes can be pharmacologically inhibited using IP3R inhibitors. However, the cAMP rise in ventricular myocytes is independent of IP3R calcium release. Our data support further investigation into the pro-arrhythmic nature of IP3-induced cAMP signaling.

## INTRODUCTION

The contraction of cardiac muscle involves the co-ordination of multiple, precisely regulated signalling pathways, summarised by the process of excitation-contraction coupling (ECC) (1). ECC is initiated by depolarisation of the sarcolemma in response to an action potential, resulting in calcium (Ca^2+^) influx across the sarcolemma, primarily through L-type Ca^2+^ channels (LTCC). In ventricular myocytes, these are located within t-tubules and positioned in close proximity to the junctional space formed between the sarcolemma and the sarcoplasmic reticulum (SR) (1) while atrial myocytes lack an extensive network of t-tubules and LTCC are primarily located in the surface membrane (2). In ventricular myocytes, ryanodine receptors (RyR) in the terminal cisternae of the SR subsequently open in response to the rise in junctional Ca^2+^ accompanying opening of LTCC, leading to further Ca^2+^ release from the SR. In atrial myocytes activation of RyR following Ca^2+^ entry via LTCC occurs by a more complex mechanism described by Blatter and colleagues as a ‘fire-diffuse-fire’ mechanism (2). Activation of Ca^2+^-induced Ca^2+^ release (CICR) in both ventricular and atrial myocytes provides an amplification of the original Ca^2+^ signal and a rise in overall cytosolic Ca^2+^ as represented by the rising phase of the cellular Ca^2+^ transient (CaT). These CaT can be recorded experimentally using intracellular Ca^2+^ sensitive dyes such as Fluo-5F. The falling phase of the CaT represents subsequent reuptake of Ca^2+^ into the SR via the sarco/endoplasmic reticulum Ca^2+^ATPase (SERCA), release from the cell across the sarcolemma, primarily via the Na^+^/Ca^2+^ exchanger (NCX), or reuptake into mitochondria (1, 3, 4). ECC provides the link between electrical excitation of cardiac tissue, in the form of an action potential, and the rise in cytosolic Ca^2+^ leading to activation of the contractile actin and myosin filaments responsible for mechanical contraction of the heart tissue (5).

The regulation of ECC in ventricular myocytes is largely influenced by (beta) β-adrenergic and muscarinic receptor signalling (1). Activation of β-adrenergic receptors (β-AR) leads to the downstream activation of adenylyl cyclase (AC), primarily AC5 and AC6 (6, 7), and elevation of cyclic adenosine monophosphate (cAMP). This cAMP binds to the catalytic subunit of protein kinase A (PKA), leading to phosphorylation and activation of downstream effectors involved in controlling inotropy including (but not limited to) LTCC (8), phospholamban (PLB) (9), RyR (10), and cardiac troponin I (cTnI) (11, 12). In addition, Ca^2+^ handling in cardiac cells can be regulated by alternative second messengers including inositol 1,4,5-trisphosphate (IP3), Ca^2+^/calmodulin-dependent protein kinase II (CaMKII), cADP-ribose (cADPR) and nicotinic acid adenine dinucleotide phosphate (NAADP) (1, 5). Specificity of cAMP/PKA signalling is achieved through compartmentalisation within specific subcellular nanodomains, primarily maintained via the action of phosphodiesterases (PDEs), which hydrolyse cAMP to AMP, and the tethering of PKA to specific targets by A-kinase anchoring proteins (AKAPs) (13–15).

IP3 is a Ca^2+^ mobilising second messenger that acts via IP3 receptors (IP3R) on the endoplasmic reticulum or SR to release Ca^2+^ in multiple cell types (16, 17). IP3 is known to play a role in Ca^2+^ handling in cardiac cells, however in comparison to ECC and the role played by RyR in CICR, the involvement of IP3 signalling in cardiac cellular Ca^2+^ handling is less well understood (18–23). Activation of G_q/11_-coupled receptors, including alpha (α)-adrenergic receptors (α-AR), angiotensin II receptor type 1 (AT1), 5-HT_2_ and endothelin-1 receptors (ET-1), leads to the cleavage of phosphatidylinositol-4,5 biphosphate (PIP2) to IP3, as well as diacylglycerol (24) and protein kinase C (PKC), via stimulation of phospholipase C (PLC) (16). In cardiomyocytes, binding of IP3 to IP3R on the SR leads to SR Ca^2+^ release (16, 17), enhancing cellular Ca^2+^ oscillations (25). However, compared to the release of SR Ca^2+^ via RyR, IP3R mediated SR Ca^2+^ release is relatively slow, and insufficient to trigger the CICR cascade (26, 27). IP3R structure is partially homologous to that of RyR, and comprises 4 identical subunits, each comprising approximately 2700 amino acids and containing an N-terminal IP3 binding sequence as well as other regulatory factors (21, 28, 29). In addition to IP3, IP3R may be regulated by cAMP/PKA as well as Ca^2+^ itself (30). In ventricular cells, IP3R is primarily located within the nuclear envelope (31), with some expression in t-tubules (32), and is involved in the coupling of nuclear Ca^2+^ signalling to gene expression (33). In atrial cells however, IP3R is expressed in the subsarcolemmal space as well as the nuclear envelope and may play a greater role in regulating Ca^2+^ handling and inotropy (19, 34). The spatial localisation of IP3Rs within cells allows IP3R mediated Ca^2+^ release to be specifically targeted to specific effectors within local signalling domains, including other IP3Rs, mitochondria, and lysosomes, as well as Ca^2+^ regulated channels and enzymes (30).

The precise role of IP3 and IP3R mediated SR Ca^2+^ release in contributing to Ca^2+^ handling in cardiomyocytes remains unclear (5, 19, 27, 35). IP3 is positively inotropic in atrial (18), and ventricular tissue (36), and positively chronotropic in the sinoatrial node (SAN) (37, 38). IP3 and IP3R are involved in controlling pacemaker and spontaneous activity of cardiomyocytes in the embryonic heart (39–41), and in the diseased heart, the re-emergence of gene expression profiles more closely replicating the embryonic condition results in IP3 having a more prominent role in these processes (21, 42, 43). Changes in the expression of IP3R have been linked with cardiac diseases, including atrial arrythmias and heat failure (18, 44, 45). Expression of IP3R type 2 (IP3R2), the most abundant IP3R isoform in cardiac tissue (18, 37), is at least 6 times greater in the atria compared to the ventricle (18) and IP3R expression is upregulated in both human patients with chronic atrial fibrillation (AF) (44) as well as canine AF models (45). Elevated Ca^2+^ and Ca^2+^ oscillations resulting from IP3R Ca^2+^ release could contribute to arrhythmogenesis (27), for example; through the stimulation of transient inward current (I*_ti_*), leading to delayed after depolarisations (DADs) (46, 47); by delaying repolarisation and increasing the likelihood of re-entry (48); or through slowing conduction (27, 49). IP3 is upregulated in patients with end stage heart failure, coinciding with a downregulation of RyR (35) and IP3R expression increases with age (19, 35, 50), an observation that parallels age-related AF disease progression from paroxysmal to persistent forms (19, 51, 52).

The inotropic and chronotropic effects of IP3 in atrial tissue are dependent upon AC and PKA activity, as shown by the inhibition of IP3 mediated changes in atrial contractility and spontaneous beating rate by the non-specific AC inhibitor MDL-12,330A and the PKA inhibitor H89 (23). These effects are independent of β-AR activation and remain in the presence of the β-AR inhibitor metoprolol (23). It has been suggested that the effects of IP3R-mediated Ca^2+^ release on cardiomyocyte contractility results from increased [Ca^2+^] within the vicinity of RyR, thereby enhancing the response of RyR to Ca^2+^ influx via LTCC (21, 53). However, the observation that changes in Ca^2+^-transient amplitude in response to IP3-producing agonists are inhibited by MDL-12,330A (MDL) and H89 implicates a role for ACs downstream of IP3R activation and independent of direct RyR sensitisation (23).

A possible explanation for the involvement of AC activity downstream of IP3R Ca^2+^ release is the direct stimulation of Ca^2+^-activated AC isoforms, AC1 or AC8, which are both expressed in close proximity to IP3R in atrial and SAN cells (23, 54–56), and are thought to influence SAN pacemaker activity through influence on the pacemaker current (*I*_f_) (57). Increases in SAN beating rate in response to activation of α-AR by phenylephrine (PE) are dependent upon AC1 activity as shown using the AC1-selective inhibitor ST034307 (56), however it remains unclear whether Ca^2+^-activated ACs are, directly or indirectly, involved in the downstream effects of IP3 signalling, or whether activation of IP3R leads to a Ca^2+^-dependent activation of AC and localised changes in cAMP within cardiomyocytes. Fluorescence resonance energy transfer (FRET)-based sensors enable real-time measurement of intracellular cAMP signalling and can be used to demonstrate compartmentalisation of cAMP/PKA signalling (14, 15, 58). Cytosolic sensors, such as EPAC-S^H187^ enable global changes in cytosolic cAMP to be measured from intact cells in real-time using fluorescent imaging (59), whereas fusion of PKA/cAMP FRET-based sensors to specific targeting domains such as AKAPs can be used to measure localised cAMP changes to determine compartmentalisation of cAMP/PKA signalling events within subcellular nanodomains (14, 15, 58, 60). Localisation of cAMP/PKA signalling, resulting from the tethering of PKA by anchoring proteins such as AKAPs (61, 62) and β-arrestins (63, 64), as well as compartmentalisation within caveolae (65, 66), accounts for the specificity of responses to cAMP. The purpose of the present study was to use a combination of FRET-based reporters, including the cytosolic sensor EPAC-S^H187^, and plasma membrane localised AKAP79-CUTie, in combination with specific IP3R inhibitors, to determine whether activation of the IP3 signalling pathway in cardiomyocytes is linked to localised changes in cAMP/PKA signals.

## MATERIALS AND METHODS

All experiments were performed in accordance with the United Kingdom Home Office guide on the Operation of Animal (Scientific Procedures) Act 1986. Biopsies of the right atrial appendage were obtained from patients undergoing on-pump cardiac surgery at the John Radcliffe Hospital (Oxford, United Kingdom). The study was approved by the Research Ethics Committee (reference no. 07/Q1607/38), and all patients gave written, informed consent.

### Cell isolation and infection

Neonatal rat atrial myocytes (NRAMs) were isolated using a modified protocol based on previous methods used to isolate neonatal rat ventricular myocytes (NRVMs) (67, 68). Hearts were isolated from 3-day-old Sprague Dawley rat pups, culled by cervical dislocation. Following dissection, atria were separated from the ventricles, transferred to 2ml Dulbecco’s Modified Eagle’s Medium (DMEM) and cut into 1-2mm^3^ pieces. Atrial myocytes were enzymatically isolated by a series of enzymatic digestions; in trypsin (1mg/ml, Merck, UK, rocked at 4°C for 2 hours) followed by collagenase (type IV, 1mg/ml, Merck, UK) as described in Burton *et al.* (67). For the collagenase digestions, the trypsin was replaced by 4ml collagenase solution and tissue was gently triturated using a plastic pipette for 1 min. The tissue plus collagenase solution was then stirred gently in a bath at 37°C for 2 mins and the first supernatant was discarded. A further 4ml collagenase solution was added to the tissue pellet and triturated for 1 min using a wide bore pipette. This solution was then stirred gently in a bath at 37°C for 2 mins and the supernatant (4ml) added to 3ml Hank’s buffered salt solution (HBSS) and stored in a 15ml centrifuge tube. This process was repeated a further 3 times to produce a total of 4 x7ml cell suspensions. Tubes were centrifuged at 2000 rpm for 8 minutes. The supernatant was then removed, and the cells were resuspended in 2ml cardiomyocyte plating media (CPM: 85% DMEM, 17% M199, 10% horse serum, 5% FBS, 1% penicillin/streptomycin) before being centrifuged at 100 g for 10 mins. All samples were then strained using a cell strainer and pooled into a single 50ml centrifuge tube. Isolated cells were pre-plated and incubated at 37°C (95% O_2_, 5% CO_2_) for 1 hour to allow separation of fibroblasts. Supernatant containing suspended atrial myocytes was then removed and cell density measured using a haemocytometer and trypan blue. Myocytes were seeded onto 24mm glass coverslips coated with 450µl laminin (40 μg/ml, Merck, UK) in 35mm 6-well plates at a density of 15,000 cells per ml in CPM. NRVMs were isolated as previously described (58). Cell media was changed every 24 hours. Cells were infected with EPAC-S^H187^ or AKAP79-CUTie second-generation adenoviral vectors after 72 hours and incubated at 37°C (14). Imaging was carried out 24-36 hours after infection.

### FRET imaging

FRET imaging was performed in an organ bath at room temperature 24 hours after infection using an inverted microscope (Olympus IX71) with a PlanApoN, 60, NA 1.42 oil immersion objective, 0.17/FN 26.5 (Olympus, UK) attached to a coolSNAP HQ^2^ camera and a DV2 optical beam splitter (MAG Biosystems, Photometrics, UK) for simultaneous recording of YFP and CFP emissions. Cells were maintained at room temperature in a modified Ringer solution (125mM NaCl, 20mM Hepes, 1mM Na_3_PO_4_, 5mM KCl, 1mM MgSO_4_, 5.5mM glucose, CaCl_2_ 1mM, pH 7.4 with NaOH). Pipetting time control experiments were conducted to ensure results are not confounded by the solvent or time. Pipetting time control consisted of the addition of either E1 on its own (for PE) or DMSO and 0.0001% pluronic F127 with 0.009% DMSO (solvent for IP3-AM) dissolved in E1. Images were acquired and processed using MetaFluor 7.1 (Meta Imaging Series, Molecular Devices). FRET ratio changes were measured as changes of the background-subtracted 480nm/545nm fluorescence (cyan/yellow) emission intensity on excitation at 430nm and expressed as *R/R_0_* where *R* is the ratio at time *t* and *R_0_* is the average ratio of the first 240 s.

### Multi-electrode Array (MEA)

For multi-electrode array (MEA) experiments, Human iPSC-derived atrial cardiomyocytes (hiPSC-ACMs) (ax2518, Axol Bioscience Ltd., UK) were cultured at 5.0×10^5^ cells/well on a 768-channel 48-well Axion Cytoview MEA 48 plate (M768-tMEA-48B, Axion Biosystems, USA) coated with Axol’s Fibronectin Coating Solution (ax0049, Axol Bioscience Ltd., UK) at 37°C in a 5% CO_2_/95% air atmosphere. Cells were cultured to day 19 after plating and all experiments were performed on day 19. Spontaneous extracellular field potentials were acquired at 37°C using a high-throughput Maestro Pro MEA system and AxIS software (v2.4, Axion Biosystems, USA). Extracellular potentials were simultaneously recorded for 16 channels per well across 48-well plates at a sampling rate of 12.5 kHz/channel. Synchronised, spontaneous beats were first observed on most electrodes across all wells from 72 hours post-thaw. For experiments, control activity both Field Action Potential (FAP) and contractility was recorded for 5 minutes before addition of DMSO, H89 (1μM) or MDL (3μM). Following addition, FAP recordings were made for 15 minutes, followed by contractility recording for 5 minutes. PE was then added at increasing concentrations from 1-30μM and FAP followed by contractility recordings were made following each subsequent addition of PE for periods of 5 mins respectively (total recording time at each concentration of PE=10 mins). Recordings were made from an additional 12 wells throughout the experimental period as an environmental control.

### Data analysis and statistics

Data are presented as mean ± SD and statistical analysis was performed using GraphPad Prism v10.1.1. Data from individual cells were averaged per plate to give n=1 and data are represented as *n*=number of averaged plates and N=number of litters used for neonatal isolations (10-14 pups per litter). For FRET experiments, data were assumed to be non-Gaussian. To compare 2 datasets a Mann-Whitney test or student’s t-test was used. Otherwise, Kruskal-Wallis followed by Dunn’s multiple comparisons, or Friedman tests were used to compare data for unpaired (Kruskal-Wallis) or paired (Friedman’s) data as appropriate. Statistical significance, when achieved, is indicated as *P < 0.05, ** P < 0.01, *** P < 0.001, **** P < 0.0001.

## RESULTS

### Stimulation of IP3 signalling using phenylephrine causes a bi-phasic rise in cytosolic cAMP in cultured NRAMs

To test the response of NRAMS expressing the cytosolic cAMP FRET sensor EPAC-S^H187^ to β-adrenergic stimulation, we used the β-AR agonist isoprenaline (ISO, 1nM). Figure 1A shows a representative time course for FRET change, representing cytosolic cAMP levels, on application of 1nM ISO to the extracellular bath solution followed by a sensor saturating concentration of Forskolin (FSK, 10μM) and 3-isobutyl-1-methylxanthine (IBMX, 100μM). Individual traces for the CFP and YFP intensity curves corresponding to the experiment shown in Figure 1A are presented in Supplementary Figure 1A. Extracellular application of 1nM ISO to NRAMS expressing EPAC-S^H187^ resulted in an increase in the mean FRET ratio (cyan/yellow) from 0.25 ± 0.01 to a peak of 0.40 ± 0.04 after 60s, representing a FRET ratio change of 44% of the saturating FSK/IBMX response (Figure 1A). Following the peak of this increase, FRET ratio decreased steadily to a plateau of 0.30 ± 0.01 within 240 s before addition of FSK and IBMX. Quantification of the FRET ratio change in response to ISO (1nM) and the saturating stimulus compared to baseline is shown in Figure 1B. In contrast, addition of the α-AR agonist PE (3μM) resulted in a bi-phasic change in cytosolic cAMP (Figure 1C and Supplementary Figure 1B), FRET ratio resulted in an initial peak (R1) corresponding to 0.38 ± 0.07 of the saturating response, followed by a second peak (R2) of 0.31 ± 0.03 from baseline (0.21 ± 0.01) representing a FRET ration change of 21 % (R1) and 12 % (R2) of the saturating response (Figure 1C). Quantification of the FRET ratio change in response to PE (3µM) and the saturating stimulus compared to baseline is shown in Figure 1D. PE was tested at a range of concentrations from 1-10μM and the maximal change in FRET ratio for R1 for these experiments is shown in Supplementary Figure 1C-E. Based on these results, a concentration of 3μM PE was used for subsequent experiments. Quantification of the FRET ratio change in response to PE (3μM, *n*=9, N=4) for R1, R2 and the saturating stimulus compared to baseline is shown in Figure 1D. Both PE FRET change responses correspond to R1 (21.20 ± 7.43%, P<0.0001, Figure 2E) and R2 (9.67 ± 4.23%, P<0.05, Figure 2F) and were significant increases when compared to a pipetting-time control (1.47 ± 0.67%, *n*=6, N=3), see supplementary Figure 1F.

**Figure 1.**
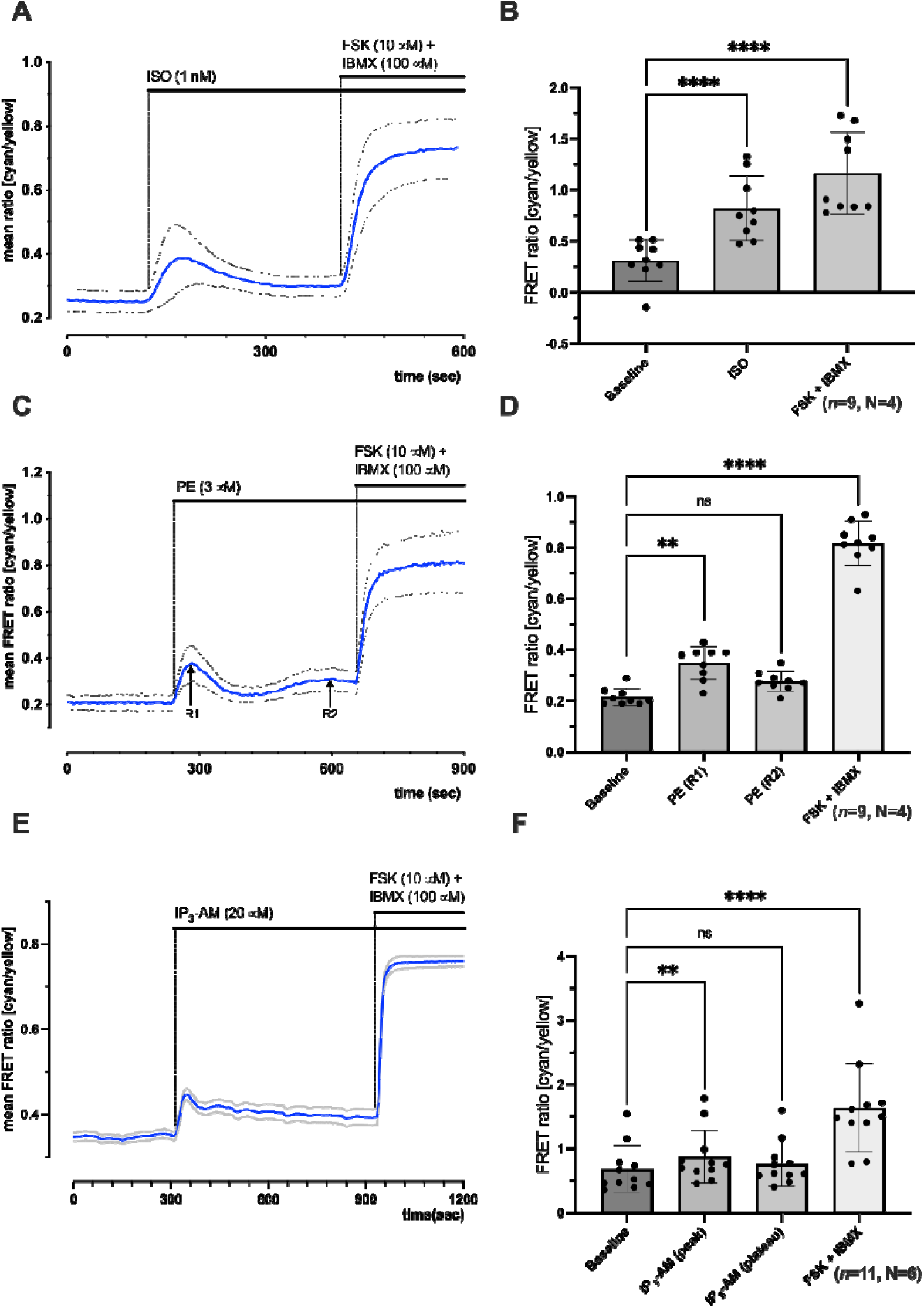
Activation of IP3 signalling pathway in NRAMs using phenylephrine results in bi-phasic increase in cytosolic cAMP as shown using the EPAC-S^H187^ FRET sensor. **A.** Representative trace from single plate (*n*=7 cells) showing time course for change in mean FRET ratio (cyan/yellow) on addition of 1nM ISO followed by 10μM FSK + 100μM IBMX. **B.** Quantification of total data showing mean FRET ratio at baseline, peak response to 1nM ISO, and saturating response to 10μM FSK + 100μM IBMX (*n*=4, N=3). **C.** Representative trace from single plate (*n*=7 cells) showing time course for change in mean FRET ratio (cyan/yellow) on addition of 3μM PE followed by 10μM FSK + 100μM IBMX. **D.** Quantification of total data showing mean FRET ratio at baseline, peak response to 3μM PE, and saturating response to 10μM FSK + 100μM IBMX (*n*=9, N=4). **E.** Representative time-course from single plate (*n*=8 cells) showing time course for change in mean FRET ratio (cyan/yellow) on addition of 20μM IP3-AM followed by 10μM FSK + 100μM IBMX. **F.** Quantification of total data showing mean FRET ratio at baseline, peak response to 20μM IP3-AM, and saturating response to 10μM FSK + 100μM IBMX (*n*=11, N=6). Representative traces and FRET ratios are represented as mean ± SD, black circles represent individual data points, ns, not significant; **P<0.01; ****P<0.0001; Friedman’s multiple comparison test.

**Figure 2.**
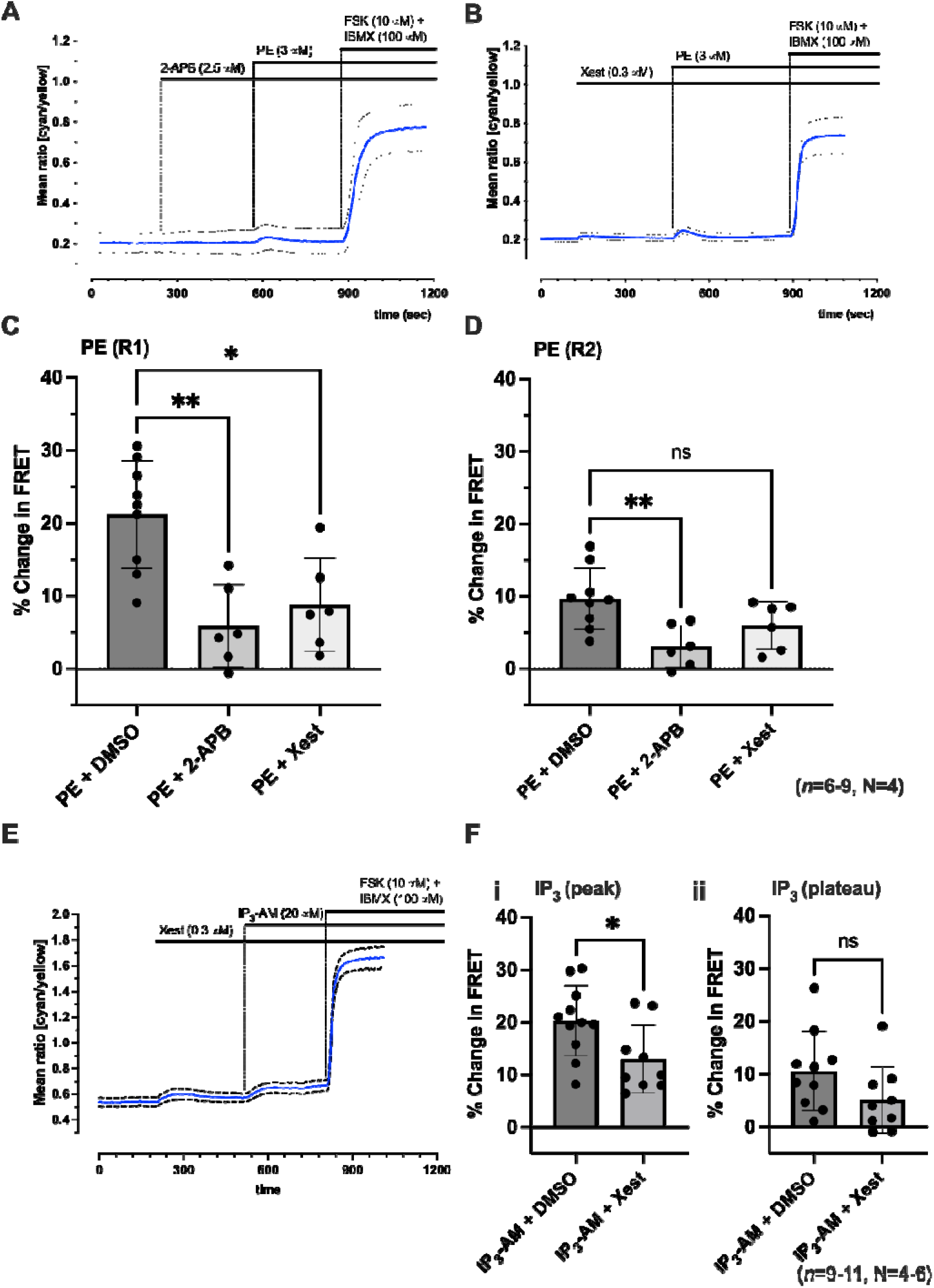
Inhibition of IP3R in NRAMs prevents rise in cytosolic cAMP in response to phenylephrine. **A** and **B.** Representative time course from single plate (**A**, *n*=6 cells; **B**, *n*=4 cells) showing time course for change in mean FRET ratio (cyan/yellow) on addition of 3μM PE followed by 10μM FSK + 100μM IBMX after initial addition of either 2.5μM 2-APB (**A**) or 0.3μM Xestospongin-C (**B**). **C.** Quantification of R1 for PE (3μM; *n*=9, N=4) alone, in the presence of 2.5μM 2-APB (*n*=6, N=4) or 0.3μM Xestospongin-C (*n*=6, N=4), expressed as a fraction of the saturating response in the presence of 10μM FSK + 100μM IBMX. **D.** Quantification of R2 for PE (3 μM; *n*=9, N=4) alone, in the presence of 2.5μM 2-APB (*n*=6, N=4) or 0.3μM Xestospongin-C (*n*=6, N=4), expressed as a fraction of the saturating response in the presence of 10μM FSK + 100μM IBMX. **E.** Representative time course from single plate (*n*=3 cells) showing time course for change in mean FRET ratio (cyan/yellow) on addition of 20μM IP3-AM followed by 10μM FSK + 100μM IBMX after initial addition of 0.3μM Xestospongin-C. **F.** Quantification of peak (i) and plateau (ii) responses for control vehicle (*n*=6, N=3) or IP3-AM (20μM; *n*=11, N=6) alone, or in the presence of 0.3μM Xestospongin-C (*n*=9, N=4), expressed as a fraction of the saturating response in the presence of 10μM FSK + 100μM IBMX. Data represented as mean ± SD, black circles represent individual data points, ns, not significant; *P<0.05; **P<0.01; unpaired student’s t-test (F) or Kruskal-Wallis followed by Dunn’s multiple comparison test (C and D).

As activation of α-adrenergic receptors (ARs) using PE may result in the activation of alternative signalling pathways via activation of PKC and DAG (16), we carried out additional experiments using the membrane permeant IP3 derivative, 2,3,6-tri-O-Butyryl-myo-IP3(1,4,5)-hexakis(acetoxymethyl)ester (IP3-AM). We found that addition of IP3-AM (20μM) to NRAMs expressing EPAC-S^H187^ produced a peak followed by a plateau phase in FRET ratio (Figure 1E). Individual traces for the CFP and YFP intensity curves corresponding to the experiment shown in Figure 1E are presented in Supplementary Figure 1G. Comparison of the quantified FRET ratio to baseline of peak and plateau following addition of IP3-AM (20μM, *n*=11, N=6) and the saturating stimulus identified a significant increase in FRET after addition of IP3-AM (P<0.01) and after addition of saturating stimuli (P<0.0001) (Figure 1F). Figure 2B shows peak FRET change corresponding to 21.20 ± 7.43% of the saturating response, followed by a plateau of 9.67 ± 4.23% of the saturating response. Both peak (21.20 ± 7.43% vs. 3.61 ± 2.04%, P<0.01) and plateau (9.67 ± 4.23% vs. 1.47 ± 2.36%, P<0.05) were significant increases when compared to a pipetting time control for IP3-AM (*n*=6, N=4) (Supplementary Figure 1H).

### Pharmacological inhibition of IP3R prevents rises in cytosolic cAMP in response to IP3-AM α-adrenergic stimulation

Figures 2A-B show representative time courses for the change in FRET recorded from NRAMs expressing EPAC-S^H187^ in response to PE (3μM) in the presence of either 2-APB (2.5μM, Figure 2A) or Xestospongin-C (0.3μM, Figure 2B). Individual traces for the CFP and YFP intensity curves corresponding to the experiment shown in Figure 2A-B are presented in Supplementary Figure 2C-D.Quantification of the effect of 2-APB and Xestospongin-C on the first (R1) and second (R2) peak, (measured as the change in FRET immediately before application of the saturation stimulus response) to PE compared to PE applied in the absence of IP3R inhibition are shown in Figures 2C (R1) and 2D (R2). Prior application of 2-APB (2.5μM) inhibited the peak change (R1) in FRET following addition of PE (3μM) from 21.20 ± 7.43% (*n*=9, N=4) to 5.89 ± 5.66% (*n*=6, N=4, *P*<0.01), whilst Xestospongin-C reduced R1 to 8.76 ± 6.43% (*n*=6, N=4, *P*<0.05) (Figure 2C). 2-APB resulted in a complete inhibition of R2 (R2 = 3.08 ± 2.89%, *n*=6, N=4, *P*<0.01 vs PE control) (Figure 2D). In the presence of Xestospongin-C, a small secondary rise in FRET was observed following R1, leading to a plateau in FRET after 300 s (Figure 2B). The amplitude of this rise (R2) was 5.96 ± 3.29% (*n*=6, N=4), which was not significantly reduced compared to the R2 measured following addition of PE alone (9.67 ± 4.23%, *n*=9, N=4, *P*>0.05) (Figures 2D).

To determine whether the changes in cytosolic cAMP recorded in response to IP3-AM and PE (Figure 1) occur downstream of IP3R activation, we used the IP3R inhibitors 2-Aminoethyl diphenylborinate (2-APB) and Xestospongin-C (Figure 2). Figures 2F shows representative time courses for the change in FRET ratio recorded from NRAMs expressing EPAC-S^H187^ in response to IP3-AM (20μM) in the presence of Xestospongin-C (0.3μM). Individual traces for the CFP and YFP intensity curves corresponding to the experiment shown in Figure 2A are presented in Supplementary Figure 2A. Quantification of the effect of Xestospongin-C on the peak and plateau (measured as the change in FRET ratio immediately before application of the saturation stimulus) response to IP3-AM compared to IP3-AM applied in the absence of IP3R inhibition is shown in Figures 2F. Prior application of Xestospongin-C (*n*=9, N=4) reduced peak to 8.76 ± 6.43% (*P*<0.05) but did not significantly reduce the amplitude of plateau 5.96 ± 3.23% (*P*>0.05), following addition of IP3-AM after DMSO control. Xestospongin C (*n*=21, N=10) alone had no effect on the cytosolic FRET signal prior to the addition of IP3-AM or PE compared to control (*n*=34, N=15) (Supplementary Figure 2B) (2.36 ± 5.74% vs. 2.41± 4.42%, P>0.9999).

### Activation of cAMP downstream of IP3R in cultured neonatal ventricular cardiomyocytes

As IP3R expression has been shown to be greater in atrial cells compared to ventricular cells (18), we were interested in determining whether cAMP changes in response to α-adrenergic stimulation would differ in NRVMs compared to NRAMs. As shown by the example trace in Figure 3A, a rise in cytosolic cAMP was observed in NRVMs expressing EPAC-S^H187^ in response to PE, with an amplitude of 14.07 ± 2.97% (*n*=10, N=6), however in contrast to the cAMP rises observed in NRAMs (Figure 1C) this rise was monophasic. The rise in cAMP observed in response to PE in NRVMs did not differ from the R1 peak observed in response to PE in NRAMs (P < 0.05, Figure 3A). In addition to PE, experiments using ISO (1nM) were also performed. Additionally, the rise in cAMP observed in response to 1nM ISO was also non-significant in NRVMs 72.26 ± 9.59% (n=8, N=6) than NRAMs 53.97 ± 14.84% (n=8, N=3) (P < 0.05, Figure 3A). The representative traces Figures Di-iii and Figures Ei-iii show the effect of PE and ISO on NRVMs in the presence of 2.5μM 2-APB (Figure 3C) and 0.3μM Xestospongin-C. Individual traces for the CFP and YFP intensity curves corresponding to the experiment shown in Figure 3D and 3E are presented in Supplementary Figure 3. In contrast to NRAMs (Figure 2), rises in cAMP in response to PE and ISO in NRVMs were not found to undergo inhibition by 2-APB or Xestospongin-C (Figure 3B and 3C).

**Figure 3.**
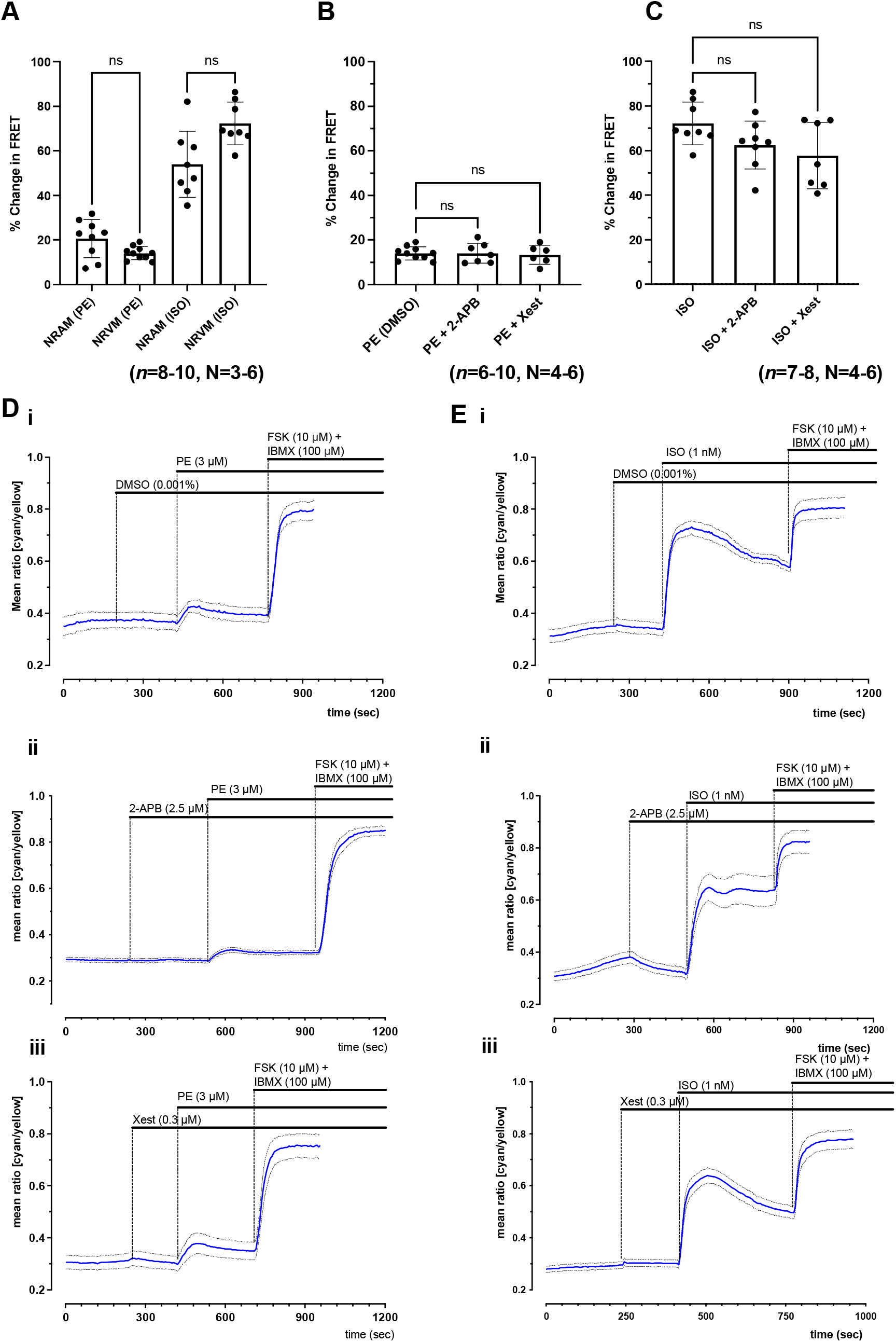
PE and ISO stimulation on cytosolic cAMP levels recorded in ventricular myocytes expressing EPAC-S^H187^ FRET sensor show that response to PE is not affected by IP3R inhibition in ventricular cells. **A.** Comparison of peak response to 3μM PE or 1nM ISO in NRAMs and NRVMs, shown as % FRET change from baseline as normalised to saturating FSK/IBMX (*n*=8-10, N=3-6). **B.** Quantification of the peak FRET change in response to 3μM PE in NRVMs in the presence of 1μl/ml DMSO, 2.5μM 2-APB or 0.3μM Xestospongin-C in NRVMs (*n*=6-10, N=4-6). **C.** Quantification of the peak FRET change in response to 1nM ISO in NRVMs in the presence of 1μl/ml DMSO, 2.5μM 2-APB or 0.3μM Xestospongin-C in NRVMs (*n*=7-8, N=4-6). **D.** Representative traces from single plates (n = 6, 14, 8 cells) showing time course for change in mean FRET ratio (cyan/yellow) on addition of 1nM ISO followed by 10μM FSK + 100μM IBMX in control conditions (i) or the presence of 2.5μM 2-APB (ii) or 0.3μM Xestospongin-C (iii) in NRVMs. **E.** Representative traces from single plates (*n*=6, 14, 8 cells) showing time course for change in mean FRET ratio (cyan/yellow) on addition of 3 μM PE followed by 10μM FSK + 100μM IBMX in control conditions (i) or the presence of 2.5μM 2-APB (ii) or 0.3μM Xestospongin-C (iii) in NRVMs. Data shown as mean ± SD, Kruskal Wallis followed by Dunn’s multiple comparison, ns=P>0.05.

### Rises in cAMP in response to α-adrenergic stimulation are localised at the plasma membrane in neonatal rat atrial cells

To test the localisation of cAMP signals in response to α-AR stimulation, we used cells expressing the plasma-membrane bound FRET sensor AKAP79-CUTie, which is targeted to the AKAP79/β-AR/AC/LTCC complex located at the internal surface of the plasmalemma facing the intracellular space (14). Figure 4A shows representative images comparing the distribution of cAMP signal of the cytosolic EPAC-S^H187^ FRET sensor and AKAP79-CUTie. The cAMP signal detected by AKAP79-CUTie was reduced compared to EPAC-S^H187^ and showed localisation at the periphery of the cell, as would be expected to coincide with AKAP79 expression, and in agreement with data reported previously (14).

**Figure 4.**
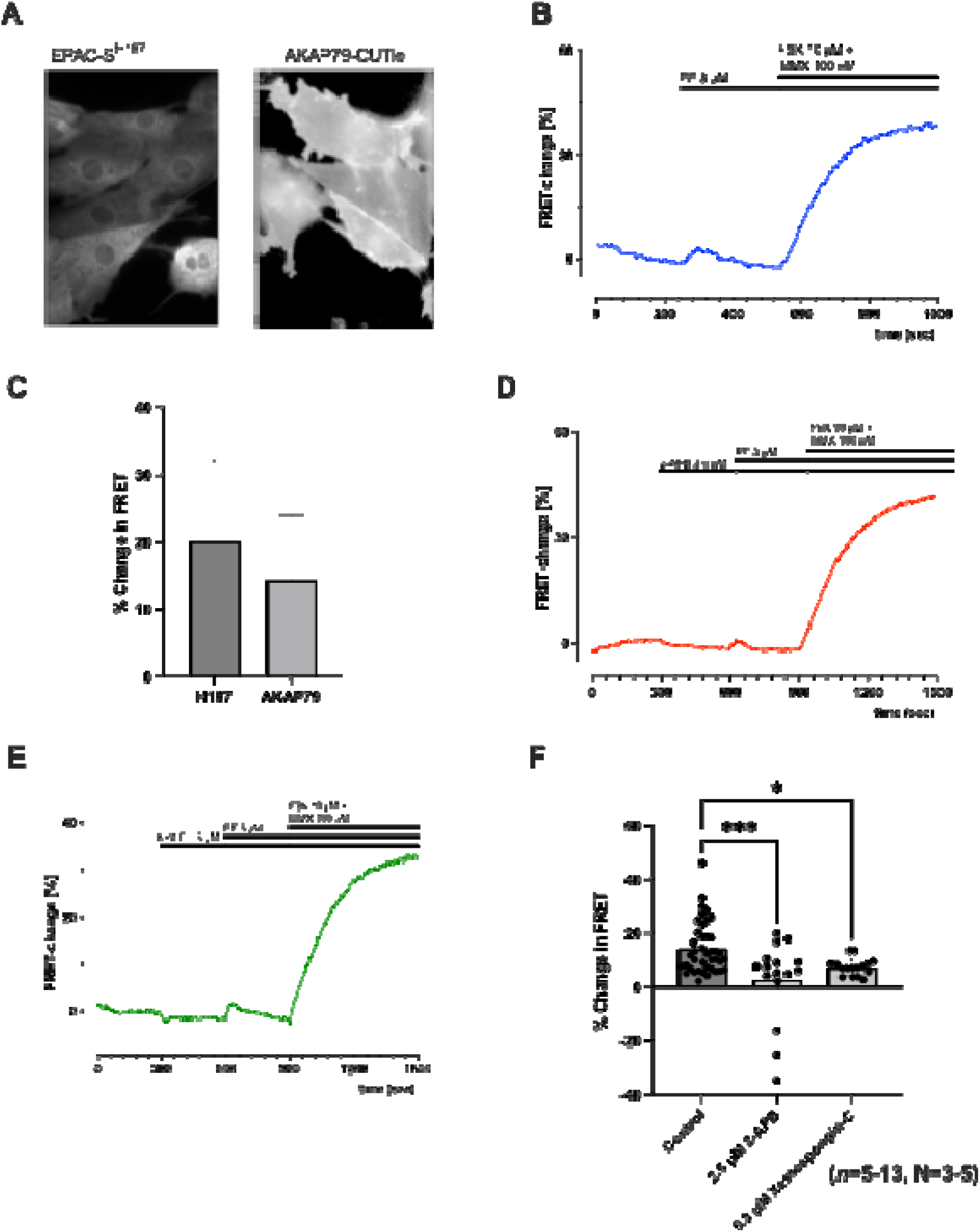
Increases in cAMP in response to IP3R activation are localised at the plasma membrane, as shown using the AKAP79-CUTie FRET sensor. **A.** Images showing representative staining in NRAM cells for EPAC-SH^187^ and AKAP79-CUTie FRET sensors. **B.** Representative trace showing time course for change in FRET ratio (yellow/cyan) recorded from cells expressing AKAP79-CUTie on addition of 3μM PE followed by 10μM FSK + 100μM IBMX (representative trace from single cell). **C.** Quantification of total data showing mean FRET ratio at baseline, peak response to 3μM PE, and saturating response to 10μM FSK + 100μM IBMX (*n*=13, N=5). **D** and **E.** Representative trace showing time course for change in FRET ratio (yellow/cyan) recorded from cells expressing AKAP79-CUTie on addition of 2.5μM 2-APB (D) or 0.3μM Xestospongin-C (E), followed by 3μM PE and 10μM FSK + 100μM IBMX (representative trace from single cell). **F.** Quantification of peak response to PE (3μM; *n*=13, N=5) alone, or in the presence of 2.5μM 2-APB (*n*=5, N=3) or 0.3μM Xestospongin-C (*n*=6, N=3), expressed as a fraction of the saturating response in the presence of 10μM FSK + 100μM IBMX. Data represented as mean ± SD, black circles represent individual data points, *P<0.05; ****P<0.0001; unpaired students t-test (C) or Kruskal-Wallis followed by Dunn’s multiple comparison (F).

Figure 4B shows a representative trace indicating the change in FRET signal from AKAP79-CUTie expressed in NRAMs on addition of PE (3μM) followed by FSK (10μM) and IBMX (100μM). Quantification and comparison of the changes of FRET for either EPAC-S^H187^ or AKAP79-CUTie expressed in NRAMs in response to 3μM PE, as a percentage of the saturating response to FSK/IBMX, are shown in Figure 4C. As shown in Figure 4F, cells expressing AKAP79-CUTie showed a 14.64 ± 2.41% increase in global FRET signal on addition of PE (P<0.05, *n*=13, N=5), indicating a rise in cAMP at the location of AKAP79/β-AR/AC/LTCC complex. Figures 4D and E show representative traces for the change in FRET signal observed in NRAMs expressing AKAP79-CUTie in response to 3μM PE in the presence of either 2-APB (2.5μM, Figure 4D) or Xestospongin C (0.3μM, Figure 4E) followed by FSK (10μM) and IBMX (100μM). 2-APB was found to cause a significant reduction in the change in FRET signal corresponding to 2.88± 15.2% (*n*=5, N=3) in response to PE compared to PE alone (Figure 4F, P<0.001). Whilst Xestospongin-C resulted in a smaller reduction in the change of FRET corresponding to 7.31 ± 3.24% (*n*=6, N=3) in response to PE, a significant reduction was still observed (P<0.05).

### Pharmacological inhibition of AC and PKA in hiPSC-ACMs inhibits the response to PE

To demonstrate the involvement of AC and PKA in the response to PE in human atrial cells we performed multi-electrode array (MEA) experiments using human-derived hiPSC-ACMs. Figure 5A shows representative averaged MEA field action potential waveforms from control (DMSO), PE (30μM), and PE (1 or 30μM) in the presence of either H89 (1μM) or MDL (3μM). Traces have been zoomed in to highlight differences in T-wave and QT interval and demonstrate consistent shortening of the Q-T interval in the presence of PE. As shown in Figure 5B, addition of PE at concentrations of 1, 3, 10 and 30μM caused a successive increase in beat rate (*n*=7), which was inhibited by H89 at high concentrations of PE (10 and 30μM PE; *n*=7; P<0.05). Addition of MDL rapidly caused all well activity to cease with increasing doses of PE (*n*=1-10).

**Figure 5.**
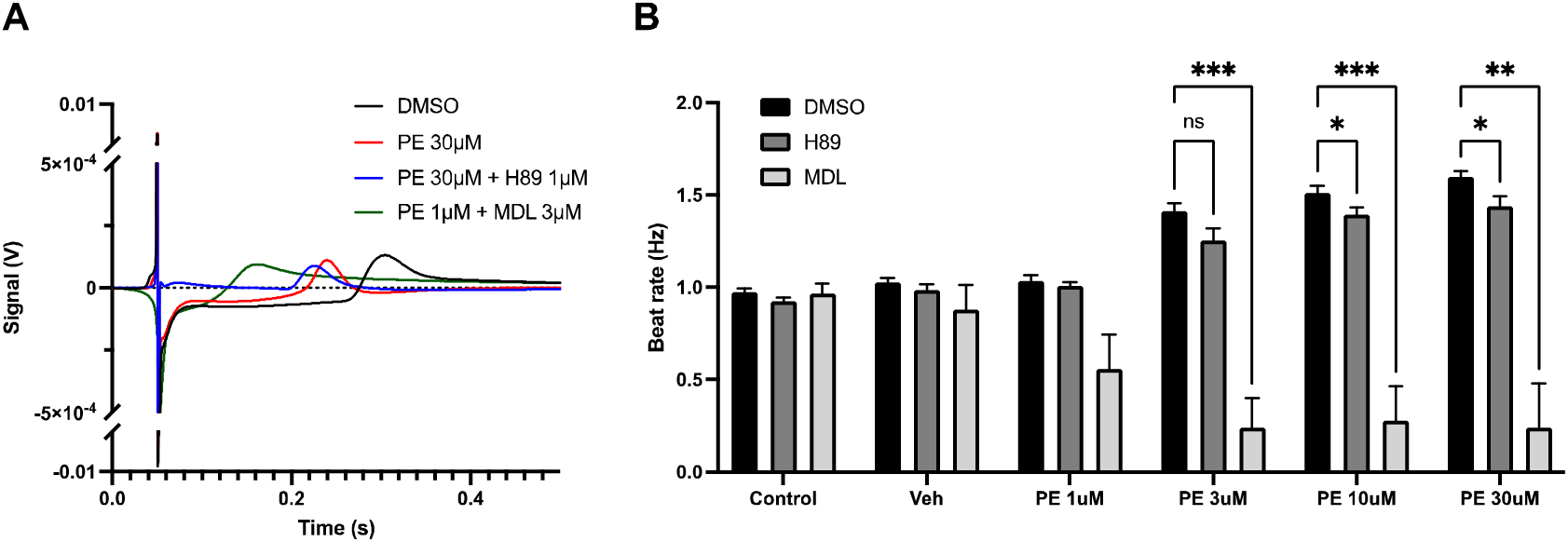
The IP3 pathway is active in human atrial iPSCs as shown using multi-electrode array. **A.** Representative averaged MEA field action potential waveforms from control (DMSO; *n*=7), PE (30μM; *n*=11), and PE (1 or 30μM) in the presence of either H89 (1μM; *n*=7) or MDL (3μM; *n*=1-10). **B.** Effect of H89 (1μM) and MDL (3μM) on the beat rate of hiPSC-ACMs in response to PE at concentrations of 1-30μM (DMSO, *n*=7; H89, *n*=7; MDL *n*=10). Data represented as mean ± SD, *P<0.05; **P<0.01; *** P<0.001; two-way repeated measures ANOVA followed by Sidak’s multiple comparisons.

## DISCUSSION

The IP3 signalling pathway has important roles to play in the control of cardiac activity, both in regulating inotropy as well as the regulation of cardiac pacemaker activity (18, 21, 23, 37). Furthermore, increasing evidence highlights the involvement of IP3 signalling in cardiac arrhythmogenesis. However, the mechanisms involved are not yet fully understood (19, 23, 34, 69).

In both atrial and ventricular cells, L-type Ca^2+^ current (*I*_Ca,L_) has been shown to undergo regulation by cytosolic Ca^2+^ (70). In ventricular cells, this regulation may be primarily modulated via CaMKII, as shown by the inhibition of I_Ca,L_ by KN-93 (70), however in atrial myocytes, the addition of the Ca^2+^ selective chelator BAPTA, following KN-93, results in a further inhibition of I_Ca,L_ not observed in ventricular myocytes, suggesting the presence of a CaMKII-independent mechanism of Ca^2+^ regulation (70). These effects of BAPTA are reversed following addition of exogenous cAMP to the cell (70). One potential mechanism that has been proposed to explain these observations, is the activation of Ca^2+^-sensitive ACs, downstream of IP3R Ca^2+^ release (23, 70). The Ca^2+^-sensitive AC isoforms AC1 and AC8 both show expression in cardiomyocytes, including atrial cells and the SAN (23, 54).

An alternative mechanism that has been proposed for linking IP3R Ca^2+^ release with effects on CaT amplitude and contractility is that Ca^2+^ released from IP3R leads to an increase in [Ca^2+^] within the immediate vicinity of RyR, which in turn, enhances the response of RyR to Ca^2+^ influx via LTCC resulting in an amplification of CICR (21, 53). In isolated atrial cells however, the enhancement of Ca^2+^ signal resulting from either pharmacological G_q/11_-coupled receptor stimulation, or direct cytosolic release of caged-IP3, is inhibited following antagonism of ACs by MDL, or PKA by H89 (23) and our data in the present study also supports the presence of this pathway in human cells as shown by the inhibition of the response to PE in hiPSC-ACMs (Figure 5). Whilst LTCCs are regulated by AC5 and AC6 (1, 71), and *I*_Ca,L_ is inhibited by MDL-12,330A (70), the direct sensitisation of RyR to Ca^2+^ released from IP3R would be expected to persist in the presence of AC inhibition. However, MDL-12,330A and H89 both result in a complete inhibition of the effects of IP3R Ca^2+^ release on CaT, indicating the involvement of AC activation downstream of IP3R activation, and before the involvement of RyR (23).

In the present study we have shown using the EPAC-S^H187^ FRET sensor that cytosolic cAMP rises in response to pharmacological activation of α-AR as well as direct application of membrane permeant IP3 (Figure 2). This change in cAMP is bi-phasic, with an initial rise (R1) followed by a secondary rise (R2) with slower kinetics (Figure 2). These rises in cAMP were inhibited following addition of the IP3R inhibitor 2-APB (Figures 2C and 2D). However, it has been reported that the effects of 2-APB on inhibiting Ca^2+^ release are related to a direct inhibitory effect on TRPC channels rather than inhibition of IP3R (64). In addition, we cannot rule out the potential of direct stimulation of TRPC following IP3R Ca^2+^ release (21). For this reason, we repeated experiments using the alternative IP3R inhibitor Xestospongin-C, and again observed an inhibition of rises in cAMP in the presence of this IP3R antagonist (Figures 2C, 2D and 2F). Potential mechanisms that could account for these changes in cAMP in response to IP3R Ca^2+^ release are discussed below:

1) cAMP may be generated by activation of AC downstream of enhanced RyR Ca^2+^ signals following IP3R Ca^2+^ release. As discussed above, this pathway appears unlikely in the models tested here due to the inhibition of changes in CaT in response to IP3 signalling in the presence of MDL-12330A and H89 (23), however additional experiments involving the inhibition of RyR would be required to confirm this.
2) Ca^2+^ release via IP3R may lead to activation of store-operated Ca^2+^ channels (SOCC), such as TRPC or Orai channels. Most likely TRPC3 as they are located on the surface membrane of SAN cells (21, 72). In addition, there is strong evidence for a functional interaction between TRPC3 (as well as TRPC6/7), IP3R, and capacitative Ca^2+^ entry (72–74). Activation of SOCC in response to IP3 could be expected to result in the downstream activation of Ca^2+^-sensitive AC1/8. In HEK293 cells, heterologous overexpression of AC1 and AC8 results in functional colocalization of AC1/8 with SOCC (75), whilst endogenously, AC8, as well as AC5/6, can be regulated by store operated Ca^2+^ entry (SOCE) (76, 77).
3) IP3R Ca^2+^ release has been shown to enhance uptake of Ca^2+^ into lysosomes (78), which themselves may act as an important intracellular Ca^2+^ store, influencing cardiac Ca^2+^ handling (79, 80). This process is itself regulated via cAMP (78, 81, 82). By boosting lysosomal [Ca^2+^], this process would be expected to enhance lysosomal Ca^2+^ release by the action of NAADP on lysosomal TPC2 channels (80).
4) Ca^2+^ released via IP3R may directly lead to the activation of Ca^2+^-sensitive AC isoforms AC1 and/or AC8 in cardiac cells. Whilst such a link has not yet been shown directly in atrial cells, in the SAN selective inhibition of Ca^2+^-sensitive AC1 using ST034307 results in a decrease in spontaneous beating rate (56), whereas chelation of Ca^2+^ using BAPTA and inhibition of ACs using MDL-12,330A reduces the hyperpolarisation activated ‘funny’ current (I_f_) (70). These data support the involvement of AC1 stimulation following IP3R Ca^2+^ release in regulation of SAN pacemaker activity (23, 37, 54, 56). As co-expression of AC1 and RyR2 in the SAN (56) appears to be comparable to that of AC8 and RyR2 in atrial cells (23), it is possible that a similar link between IP3R Ca^2+^ release and activation of AC8 is present in atrial cardiomyocytes. Furthermore, overexpression of AC8 in mouse cardiac cells has been shown to result in increased contractility, independent of *I*_Ca,L_, demonstrating that endogenous AC8 may have the ability to influence cardiac contractility via a mechanism independent of AC5/6 (83).

The biphasic pattern of cAMP generation observed in Figure 1 in response to PE likely results from the combination of more than one of the pathways discussed above. Interestingly, local changes in cAMP at the cell membrane, identified using the FRET sensor AKAP79-CUTie, only showed an initial, fast (R1) rise in cAMP (Figure 4). Whilst it is possible that a secondary rise was undetected due to the low signal generated using AKAP79-CUTie, another explanation is that the R1 rise results primarily from the activation of ACs located within the region of the plasma membrane. In contrast, the secondary R2 cAMP rise is the result of a slower cytosolic cAMP release. Such a delayed, cytosolic response could be expected to occur for example if this rise occurred downstream of lysosomal Ca^2+^ release, or secondary to TRPC or Orai channel activation. These studies highlight the need for further investigation into the link between IP3R mediated SR Ca^2+^ release and lysosomal Ca^2+^ release via TPC2 in response to NAADP as well as the potential role of SOCC and SOCE in atrial IP3 signalling.

Our data comparing the rise in cytosolic cAMP in response to PE in atrial cells with that observed in ventricular cells highlighted the absence of the secondary R2 rise in ventricular cells (Figure 3A). Furthermore, the cAMP rise observed in ventricular cells did not undergo inhibition in the presence of either 2-APB or Xestospongin-C (Figure 3E), indicating that this rise likely occurs independently of IP3R Ca^2+^ release. These data support the presence of an IP3R dependent pathway of cAMP activation that is present in atrial myocytes but absent or reduced in ventricular myocytes.

Several FRET-based sensors rely on the binding of AKAPs, principally AKAP79, for targeting subcellular domains (15). AKAP79 binds to the dimerization/docking (D/D) domain of PKA RII subunits (84) and is localised to the plasma membrane through binding with PIP2 (85). In addition, AKAP79 can interact with PKC (86) and Ca^2+^-dependent protein phosphatase 2B (PP2B) (15, 87). In cardiomyocytes, AKAP79 forms a complex with β-AR, AC5/6, and LTCC (15, 88). One potential drawback of the use of AKAP79-targeted FRET sensors is the possibility that these sensors may themselves impact the kinetics and amplitude of the local changes in cAMP. However the AKAP-79-CUTie FRET sensor has been shown to avoid these issues (15). Using AKAP79-CUTie, we observed changes in FRET within the region of the plasma membrane in response to IP3R activation via potentiation of G_q/11_-coupled receptors, using phenylephrine. AKAP79-CUTie has been previously used to identify cAMP nano-domain signalling within the region of β-AR (14), and given the association of AKAP79 with AC5/6, we cannot rule out the possibility that the cAMP changes we observed resulted from the action of these AC isoforms. AC5/6 are inhibited by Ca^2+^ (83). However, our experiments using 2-APB and Xestospongin C (Figure 4) demonstrate that the changes we observed in cAMP using AKAP79-CUTie are dependent upon either IP3R Ca^2+^ release or SOCC. AC8 would therefore be a more likely candidate for this rise in cAMP, as we know that *I*_Ca,L_ in atrial cardiomyocytes may also be regulated by Ca^2+^ dependent ACs (70), and that AC8 is expressed in this region of the plasma membrane (23, 56).

Although I_Ca,L_ and RyR2 are strongly suspected as the downstream targets of IP3-cAMP signals, the direct phosphorylation and activation by PKA in response to IP3 of these or any other target proteins has yet to be shown. Moreover, a direct role for IP3-cAMP signals in the reported pro-arrhythmic effects of IP3 signalling in atria has yet to be proven, although it appears likely if RyR2 and I_Ca,L_ are phosphorylated by PKA following IP3 signalling. Although the amplification of the CaT in atrial myocytes via IP3 is PKA dependant (Capel et al. 2021), this does not mean IP_3_ cAMP signals cannot have alternative pro-arrhythmic effects on atrial myocytes via other cAMP sensitive targets such as HCN, EPAC1/2, POP-EYE domain containing proteins and phosphodiesterase 10. Additionally, the effects of cAMP hydrolysing phosphodiesterases that may regulate the IP3-cAMP remains unknown.

As a first step towards exploring the translational potential of our cAMP and IP3 signalling findings, we attempted to perform western blots on right atrial appendage tissue samples from AF patients to look for changes in AC expression. However, this was unsuccessful as the antibodies tested failed to work on human tissue. In order not to lose our precious samples, we opted to perform data-independent acquisition parallel accumulation-serial fragmentation (DIA-PASEF) with label free protein quantification, a much more sensitive method for measuring low abundance proteins as it improves depth and quantitation of detected proteins. Through this we identified the presence of AC5 and 6 and multiple components of the IP3 signalling cascade (See supplementary Figure 4C). These data are available on PRIDE (Project Name: Activation of IP3R in atrial cardiomyocytes leads to generation of cytosolic cAMP as shown using fluorescence resonance energy transfer; Project accession: PXD042391). None of these proteins’ abundance differed significantly between AF tissue samples and control, supporting the concept that IP3 signalling is a viable target in AF patients (Supplementary Figure 4C and Supplementary Table 1). However, atrial tissue contains a range of different cell types from atrial myocytes to vascular smooth muscle cells, endothelial cells, neurons, immune cells and fibroblasts meaning changes in the expression of IP3 signalling components and AC in specific cell types within the atria in AF cannot be ruled out.

In the current paper, we have shown for the first time, using FRET-based sensors, that cytosolic cAMP levels are increased in atrial myocytes following α-AR activation (Figure 1). In atrial cells, these changes in cAMP can be inhibited by either 2-APB or Xestospongin-C, confirming that cAMP is generated subsequently to IP3R activation in atrial cardiomyocytes. Whether this rise in cAMP occurs as the result of the direct influence of Ca^2+^ release from IP3R on Ca^2+^-sensitive AC1/8, or via an intermediate effect, such as activation of SOCE, remains to be determined. Conclusive identification of the targets of IP3-cAMP signals, any pro-arrhythmic effects, and the cAMP phosphodiesterases that regulate IP3-cAMP signals in atrial myocytes may identify novel targets for the modulation of pro-arrhythmic atrial IP3 signalling.

## Supplementary Methods

Tissue sample preparation for proteomics: The right atrial appendage tissue of SR and AF were individually homogenized in a bead homogenizer in RIPA buffer and were reduced using 4X Lithium Dodecyl Sulfate (LDS) buffer and beta-mercaptoethanol. Tissue samples were analysed by LC-MS/MS using a Dionex Ultimate 3000 (Thermo Scientific) coupled to a timsTOF Pro (Bruker, (24)) using a 75μm x 150mm C18 column with 1.6μm particles (IonOpticks) at a flow rate of 400 nL/min. A 17-minute linear gradient from 2% buffer B to 30% buffer B (A: 0.1% formic acid in water. B: 0.1% formic acid in acetonitrile) was used.

Performing Mass Spec: The TimsTOF Flex (Bruker) was operated in PASEF mode using Compass Hystar 5.0.36.0. Settings for the 11 samples per day method were as follows: Mass Range 100 to 1700m/z, 1/K0 Start 0.6 V·s/cm^2^ End 1.6 V·s/cm^2^, Ramp time 110.1ms, Lock Duty Cycle to 100%, Capillary Voltage 1600V, Dry Gas 3 l/min, Dry Temp 180°C, PASEF settings: 10 MS/MS scans (total cycle time 1.27sec), charge range 0-5, active exclusion for 0.4 min, Scheduling Target intensity 10000, Intensity threshold 2500, CID collision energy 42eV. Settings for the 50 and 180 samples per day method were as follows: Mass Range 100 to 1700m/z, 1/K0 Start 0.85 V·s/cm^2^ End 1.3 V·s/cm^2^, Ramp time 100ms, Lock Duty Cycle to 100%, Capillary Voltage 1600V, Dry Gas 3 l/min, Dry Temp 180°C, PASEF settings: 4 MS/MS scans (total cycle time 0.53 sec), charge range 0-5, active exclusion for 0.4 min, Scheduling Target intensity 24000, Intensity threshold 2000, CID collision energy 42eV.

Analysis and statistics on mass spec data: The raw protein intensities of individual SR vs AF group samples were Log transformed and normalised by median subtraction to identify the significantly regulated proteins of interest in AF. The violin plots were created with the InstantClue omics tool version, win-0.12.1 (89). The protein intensities of the AF vs SR groups were log-transformed and normalized using the Z score. The kernel density estimation of protein intensities for the distribution between groups was represented using a gradient of colour code, red to blue.

## SUPPLEMENTAL MATERIAL

**Supplementary Figure 1 (related to Figure 2):**
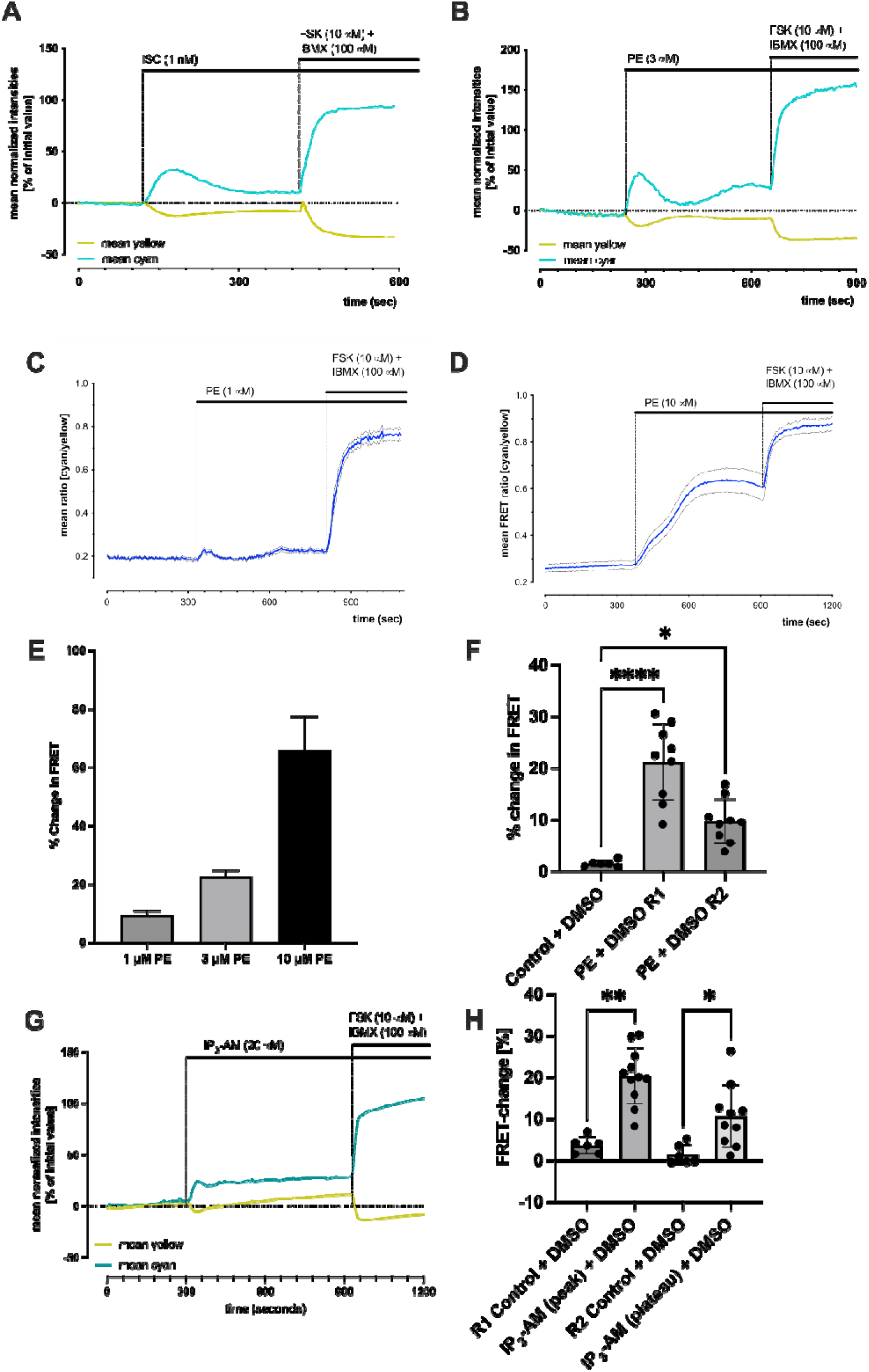
**A.** Representative example single cell trace of CFP/YFP changes in FRET signal in response to 1nM Isoprenaline in NRAMs loaded with EPAC-S^H187^. **B.** Representative example single cell trace of CFP/YFP changes in FRET signal in response to 3 μM PE in NRAMs infected with EPAC-S^H187^. **C.** Representative example single well trace of the effect of 1 μM PE on cAMP levels in NRAMs loaded with EPAC-S^H187^. **D.** Representative example of a single well trace of the effect of 10μM PE on cAMP levels in NRAMs loaded with EPAC-S^H187^. **C-D.** Solid line = mean ratio of all viable cells in well. Dotted line=SD of ratio of all viable cells in well. **E.** Comparison of the peak R1 response to 1-10μM PE on cAMP levels in NRAMs loaded with EPAC-S^H187^. Data presented as mean ± SEM. **F.** Quantification of peak changes in response to 3μM PE R1 and R2 (*n*=9, N=4) vs. time and pipette controls (*n*=6, N=3). **G.** Representative example trace of CFP/YFP changes in FRET signal in response to 20 μM IP3-AM. **H.** Quantification of peak changes in response to 20μM IP3-AM peak and plateau (*n*=11, N=6) vs. time/pipette controls (*n*=6, N=3). In **F** and **H** each block dot represents the mean reading from an experiment in a single well. In **F** and **H** data is presented as mean ± SD. In **F** and **H** data statistically tested using Kruskal-Wallis followed by Dunn’s multiple comparison. *P<0.05, **P<0.01, ***P<0.001, ****P<0.0001.

**Supplementary Figure 2 (related to Figure 2):**
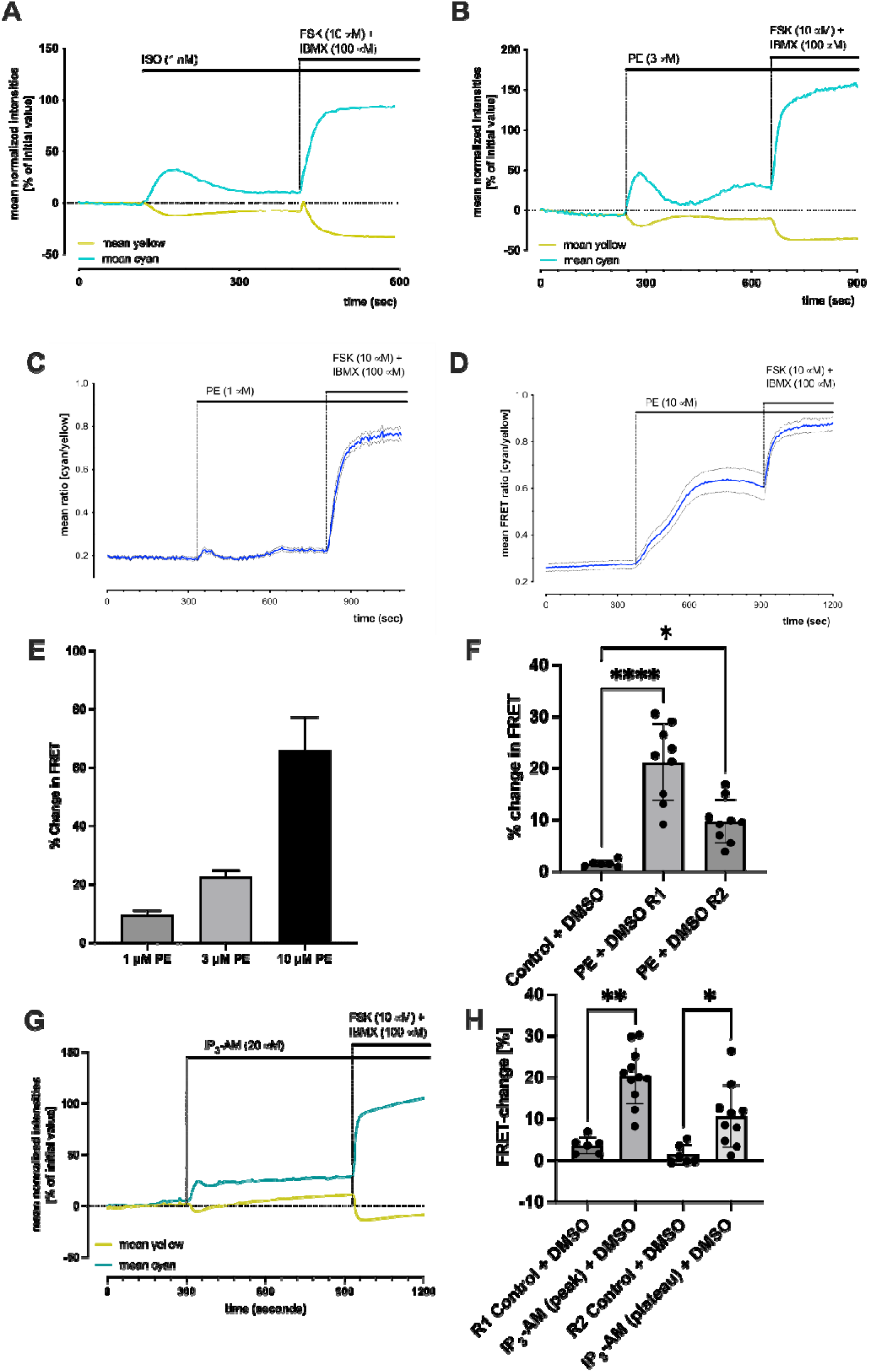
**A.** Representative example single cell trace of CFP/YFP ratio in EPAC-S^H187^ FRET signal in response to 20µM IP3-AM in the presence of 0.3µM Xestospongin C in NRAMs. **B.** Effect of 2.5µM of 2-APB and 0.3µM of Xestospongin-C on the baseline cytosolic [cAMP]. Each block dot represents the mean reading from an experiment in a single well. **C-D.** Representative example single cell trace of CFP/YFP changes in EPAC-S^H187^ FRET signal in response to 3µM PE in the presence of IP3R inhibitors 2.5µM 2-APB and 0.3µM Xestospongin-C respectively. Data are presented as mean ± SD. Data statistically tested using Kruskal-Wallis followed by Dunn’s multiple comparison. ns= non-significant. ***P<0.001, ****P<0.0001. n number = number of wells.

**Supplementary Figure 3 (related to Figure 3):**
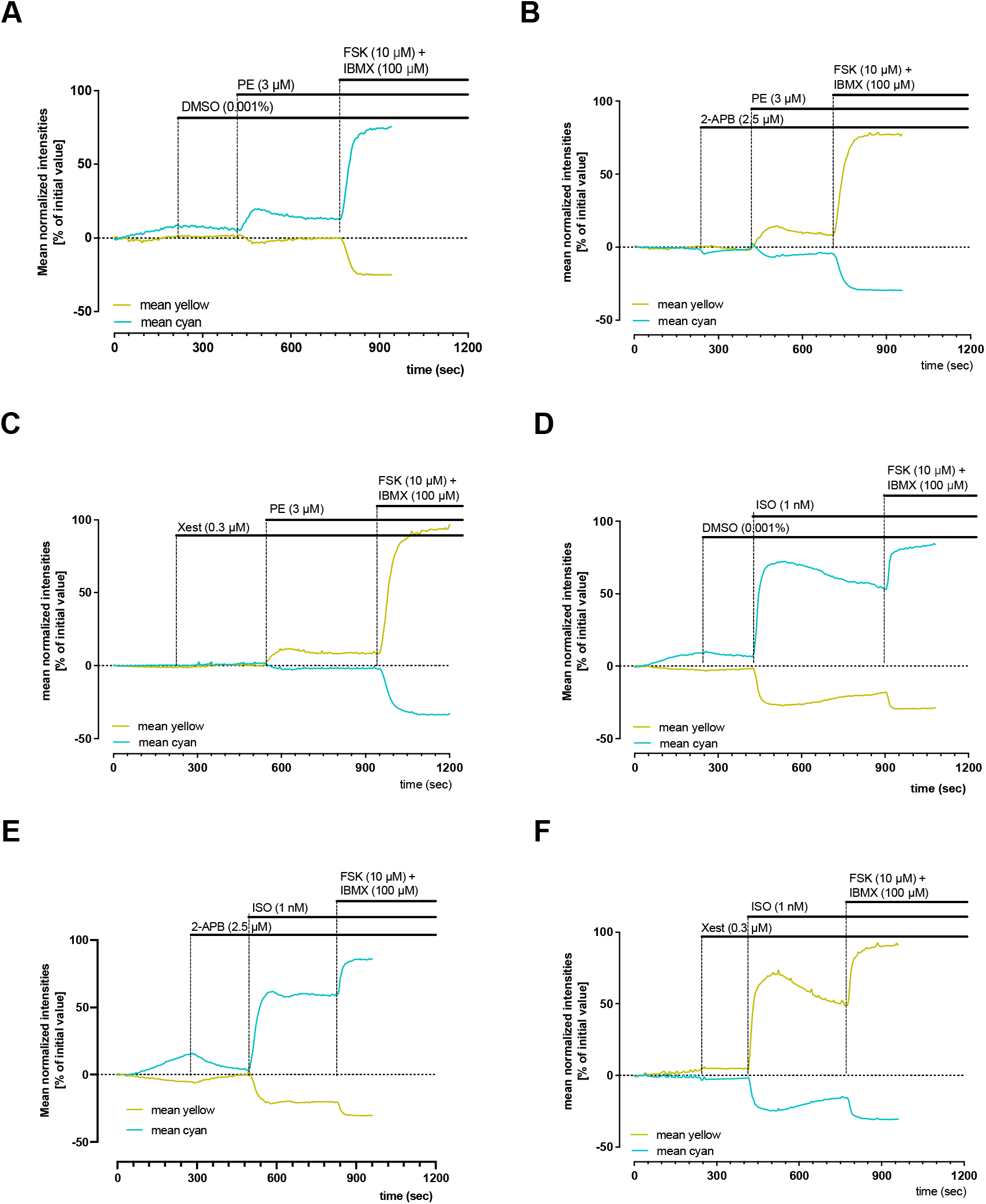
**A-C.** CFP/YFP changes in FRET signal in response to 1 nM ISO in the presence of DMSO (**A**) and IP3R inhibitors 2.5µM of 2-APB (**B**) or 0.3µM of Xestospongin-C (**C**) (*n*=6, 14, 8 cells) in NRVMs. **D-F.** CFP/YFP changes in FRET signal in response to PE in the presence of DMSO (**D**) and IP3R inhibitors 2.5µM of 2-APB (**E**) or 0.3µM of Xestospongin-C (**F**) (*n*=7, 7, 6 cells) in NRVMs.

**Supplementary Figure 4:**
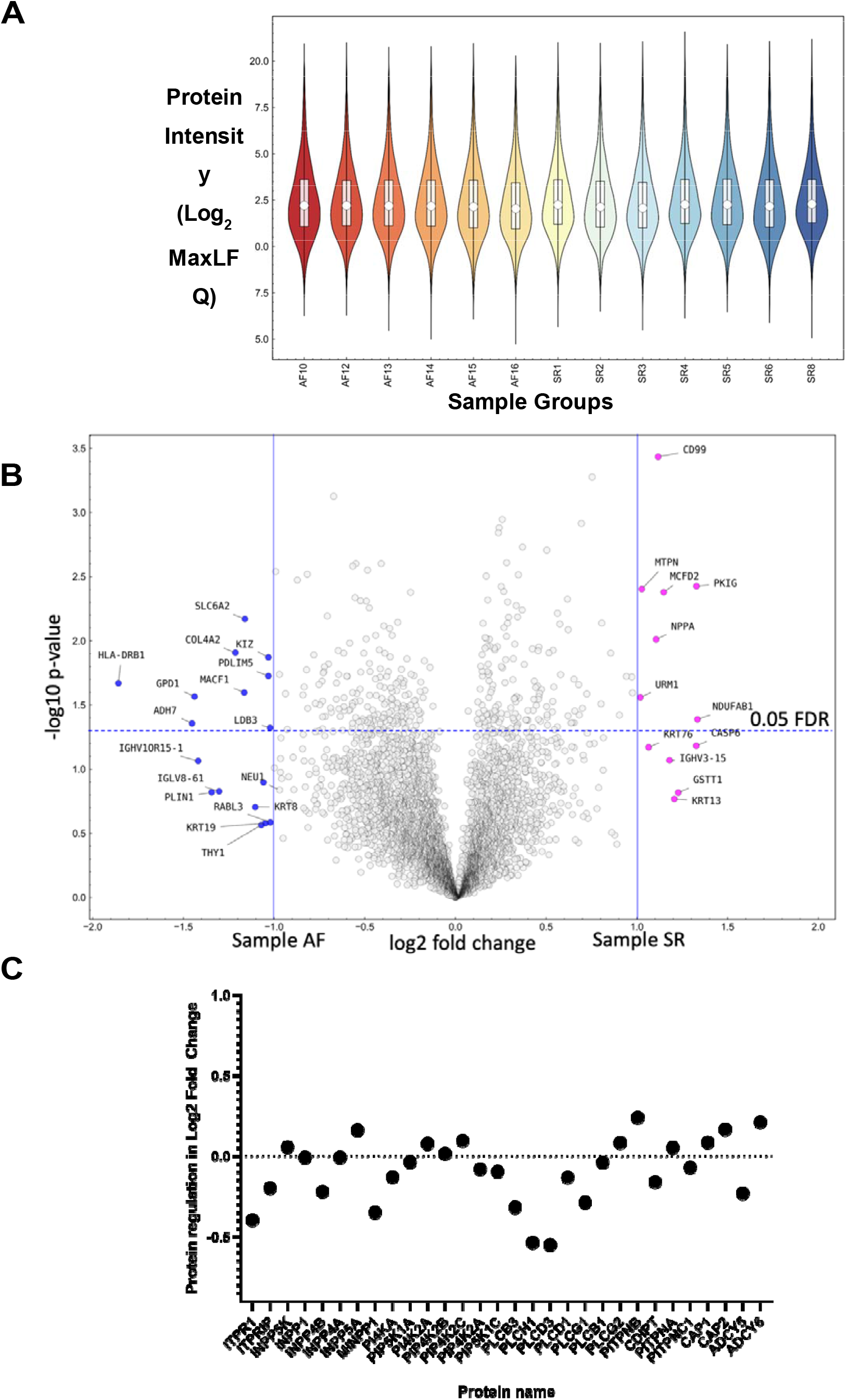
**A.** Violin Plot of protein intensity preparations in human samples from atrial fibrillation patient (N=6) and patient with a normal sinus rhythm (N=7). **B.** Volcano Plot of differential gene expression of human samples from atrial fibrillation patient (N=6) and patient with a normal sinus rhythm (N=7), MaxLFQ is a generic method for label-free quantification, proteins with fold changes greater than 2 are highlighted, Instant Clue v0.12.1. **C.** Expression of proteins of interest specifically related to IP3 signalling including ACs, phosphodiesterases from human samples from atrial fibrillation patient (N=6) normalised to patient with a normal sinus rhythm (N=7) shown in a protein regulation graph.

**Supplementary table 1:** Volcano Plot excel output table (related to supplementary figure 4B, student T-test). Used to determine significant data points in a data set with a permutation-based FDR, X-axis column displays: log2 fold changes and Y-axis column displays: -log10 p values. Column headers are gene/protein description and columns labelled with the sample IDs are showing the mass spec signal intensities in log2, normalized by MaxLFQ (90).

## ACKNOWLEDGEMENTS

This study was supported by a British Heart Foundation Project grant (PG/18/4/33521). SJB is a post-doctoral scientist funded by the British Heart Foundation (PG/18/4/33521). RABB is funded by a Sir Henry Dale Wellcome Trust and Royal Society Fellowship (109371/Z/15/Z) and holds a Senior Research Fellowship at Linacre College, Oxford. RAC is a post-doctoral scientist funded by the Wellcome Trust and Royal Society (109371/Z/15/Z). MZ is supported by the British Heart Foundation (RG/17/6/32944). TA received funding from the Returners Carers Fund (PI RABB), Medical Science Division, University of Oxford, the Nuffield Benefaction for Medicine and the Wellcome Institutional Strategic Support Fund (ISSF), University of Oxford. EA received funding from the Returners Carers Fund (PI RAC), University of Oxford. RABB acknowledges support from the Covid-19 Rebuilding Research Momentum Fund (CRRMF) Oxford Funds and Returning Carers Funding (Oxford). SDB and RA were employees of Axol Bioscience Ltd. (Easter Bush, UK).

## DISCLAMERS

None

## AUTHOR CONTRIBUTIONS

RABB, DT conceived the research. MZ supervised all FRET experimentation and design of probes. All authors contributed intellectually to the study. SJB and RABB designed the study. SJB, EA, and MJR carried out animal dissections and cell isolations. SDB and RA carried out multi-electrode array experiments. SJB and EA performed Iso and PE FRET experiments. MJR performed IP3-AM experiments and PE time/pipetting control FRET experiments. SJB wrote the first draft and EA and MJR contributed to subsequent drafting of the manuscript. TA and JS performed Western Blots on human tissue samples under BC human ethics. TA, RF and SH performed DIA-PASEF mass spectrometry experiments. All authors have contributed to the content and refinement of the manuscript.

